# Truly the best of both worlds: merging lineage-specific and universal baiting kits to maximize phylogenomic inference

**DOI:** 10.1101/2023.11.16.567445

**Authors:** Luiz Henrique M. Fonseca, Pieter Asselman, Katherine R. Goodrich, Francis J. Nge, Vincent Soulé, Kathryn Mercier, Thomas L. P. Couvreur, Lars W. Chatrou

**Author notes:** Authors for Correspondence: LHMF < >; LWC < >.

## Abstract

**PREMISE:** The development of RNA baiting kits for reduced representation approaches of genomic sequencing is popularized, with universal and clade-specific kits for flowering plants available. Here, we provided an updated version of the Annonaceae bait kit targeting 799 low copy genes, known as Annonaceae799.

**METHODS:** This new version of the kit combines the original 469 genes from the previous version of the Annonaceae kit with 334 genes from the universal Angiosperms353 kit. We also compared the results obtained using the Original Angiosperms353 kit with our custom approach. Parsimony informative sites (pis) were evaluated for all genes and combined matrices.

**RESULTS:** The new version of the kit has extremely high rates of gene recovery. On average, 796 genes were recovered per sample, and 777.5 genes recovered with at least 50% of their size. Off-target reads were also obtained. Evaluating size, the proportion of on- and off-target regions, and the number of pis, the genes from the Angiosperms353 usually outperform the genes from the original Annonaceae bait kit.

**DISCUSSION:** The results obtained show that the new sequences from the Angiosperms353 aggregate variable and putative relevant bases for future studies on species-level phylogenomics, and within species studies. The merging of kits also creates a link between projects and makes available new genes for phylogenetic and populational studies.

---

Target enrichment methods allow for the sequencing of hundreds of independent low-copy loci from the nuclear genome and cytoplasmic regions (Mamanova et al., 2010; Cronn et al., 2012; Weitemier et al., 2014). This wealth of molecular data can be applied in phylogenetic and population studies of non-model organisms, opening a new era for plant systematics and species delimitation, among other fields (Soltis et al., 2013; McKain et al., 2018; Anderman et al., 2020). Strategies of genome reduction that incorporate hundreds of thousands to millions of base pairs are allowing the inference of robust phylogenetic hypotheses incorporating the effects of diverse processes, such as incomplete lineage sorting (Pamilo and Nei, 1988; Maddison, 1997), introgression (Nakhleh, 2013), or accounting for the simultaneous effects of both (Nakhleh, 2013). Independent loci such as those from the nuclear genome are critical for species tree inference under a multispecies coalescent (MSC) model, or a multispecies network coalescent (MSNC) model when reticulation is also considered (reviewed in Wen and Nakhleh, 2018). The use of target enrichment methods can also be extended to population studies with the possibility of orthologous gene variants to be separated (phased) and single nucleotide polymorphism to be assessed (Nauheimer et al, 2021; Jiménez-Mena et al., 2022).

Gene capture techniques have been quite successful in multiple plant studies at diverse levels. Universal kits targeting all angiosperms are available (Buddenhagen et al., 2016; Johnson et al., 2019; Waycott et al., 2021), as well as customized bait kits designed for plant families such as Annonaceae (Couvreur et al., 2019), Apocynaceae (Weitemier et al., 2014), Asteraceae (Mandel et al., 2014), Bignoniaceae (Fonseca et al., 2023), Gesneriaceae (Ogutcen et al., 2021), Ochnaceae (Schneider et al., 2020) and Orchidaceae (Eserman et al., 2021); or more inclusive clades such as *Begonia* L. (Begoniaceae; Michael et al., 2022), *Buddleja* L. (Scrophulariaceae; Chau et al., 2018), *Dioscorea* L. (Dioscoreaceae; Soto Gomez et al., 2019), and *Euphorbia* L. (Euphorbiaceae; Villaverde et al., 2018). Even species-specific sets have been designed for population genetics and breeding (e.g., barley, *Hordeum vulgare* L.; Hill et al., 2019). Attempts to compare customized and universal baiting kits at shallow levels of the phylogeny often conclude that the performance of both kits is similar for phylogenetic studies (e.g., Larridon et al., 2020; Shah et al., 2021). Rather than comparing, a fruitful approach is the merging of both universal and specific gene sets in a single bait kit (Hendricks et al., 2021). By doing so, it is possible to combine the versatility of universal and the efficiency of specific bait kits in a single framework, making available as many genes as possible for phylogenetic or population studies (Mandel et al., 2014; Eserman et al., 2021; Fonseca et al., 2023).

A bait kit designed specifically for the plant family Annonaceae is available, targeting up to 469 exons (genes) and 364,653 bp (Couvreur et al., 2019). This baiting kit was designed considering the molecular variability across the whole family and was tested both at the family level (65 different genera) and the species level in the Piptostigmateae tribe (29 species, and multiple individuals) with positive results (Couvreur et al., 2019). Several recent studies have since confirmed the utility of the kit to infer species level relationships in Annonaceae (Dagallier et al., 2023; Martínez-Velarde et al., 2023; Lopes et al., in press). However, the lack of overlap between the widely used Angiosperms353 bait kit (Johnson et al., 2019), and the customized baiting kit (Couvreur et al., 2019) limits the reuse of the data generated for Annonaceae and the contribution to ongoing efforts to assemble the plant tree of life (Baker et al., 2021). The baits designed using the Angiosperms353 gene set in combination with clade specific species as references also tend to improve enrichment, putatively yielding more reads for gene assembly compared to the Original Angiosperm353 kit (Baker et al., 2021).

In this study, we address these limitations by generating an Annonaceae-specific bait set combining the exons of the 469 genes selected by Couvreur et al. (2019), with single-copy genes from the Angiosperms353 panel (Johnson et al., 2019) to build a set of 799 putatively single-copy nuclear genes (803 genes were tackled in total, with four genes failing), which we hypothesize will provide a scope for more robust estimates of relationships with greater support. We call this new baiting kit Annonaceae799. We tested the utility of this new bait kit using *Asimina* Adans., the only non-tropical genus of Annonaceae. The resolution of species-level phylogenetic relationships in this genus has proven to be problematic, especially if we consider the accounts of putative gene flow (and introgression) between species of *Asimina* and the formation of morphological intermediates treated as hybrids (Kral, 1960). Hundreds of independent nuclear genes were evaluated here regarding their variability and the presence of paralogs. Multiple individuals of *Asimina triloba* (L.) Dunal were also compared, showing the utility of the gene set within species. We also provide a detailed comparison between results obtained through the new Custom Angiosperms353 baiting set designed here with the original kit (Johnson et al., 2019).

## METHODS

### Gene selection

The original alignments from the Angiosperms353 panel (available at: https://github.com/mossmatters/Angiosperms353) were used in this study as sources of reference sequences for bait design. Sequences of Annonaceae transcriptomes from the 1KP project (Leebens-Mack et al., 2019) were kept (i.e., *Annona muricata* L. [YZRI], *Uvaria macrocarpa* Champ. ex Benth. [PSJT]), while species outside the family were excluded. For 312 genes, at least one sequence of Annonaceae was available and used as reference. For the remaining 22 genes, we used *Eupomatia bennettii* F.Muell. (DHPO) as reference. Annonaceae and Eupomatiaceae are sister families, justifying our choice (Chatrou et al., 2012). In total, exons from 803 genes were initially combined. These included the 469 genes selected by Couvreur et al. (2019) plus 334 single copy genes from the Angiosperms353 panel (Johnson et al., 2019). All genes were evaluated regarding the number of copies using the genome of *Liriodendron chinensis* (Hemsl.) Sarg. (Magnoliaceae; Chen et al., 2019; GenBank: PRJNA418360) as the closest genome available and BLAT (Kent, 2002). Duplicates were searched using the program CD-HIT-EST 4.5 (with –c 0.9) (Fu et al., 2012). Of the original 803 genes, four were flagged by CD-HIT-EST for having similar or identical copies, in which case the longest copy available was kept. The baits were designed using 799 genes covering 698,154 bp. Using the original alignments from the 1KP project (Johnson et al., 2019; Leebens-Mack et al., 2019) and keeping only sequences from Annonaceae or closely related species, we were able to save thousands of baits in our final set, enabling the merger of both Angiosperms353 and Annonaceae bait kits in a single set.

### Sampling

We sampled 22 specimens of *Asimina* and outgroups, including all 11 known species within the focus genera. As taxonomic reference for *Asimina*, we used Kral (1960) and the Flora of North America (http://floranorthamerica.org/). We also included the two species of *Deeringothamnus*, previously recognized as a separate genus based on morphology (Wilbur, 1970), but recently recovered as nested within *Asimina* in a molecular phylogenetic study (Li et al., 2017). For *A. triloba*, seven specimens from different localities were included in our study. As outgroups, we selected the closely related *Annona glabra* L., *Annona scandens* Diels, *Diclinanona calycina* (Diels) R.E.Fr., and *Goniothalamus laoticus* (Finet & Gagnep.) Bân (Tables 1; S1) (Erkens et al., 2014). For most of the species, leaves were collected in the field in 2006, 2021, and 2022 and dried in silica gel. To complement the sampling, herbarium material was also used as source of leaf tissue (herbaria FLAS and L; Table S1).

### DNA extraction and library construction

Total DNA was extracted using the 2x CTAB method (Doyle & Doyle, 1987). The step of cell lysis step was carried out overnight at 60° C. Total DNA extractions were evaluated by means of integrity using agarose gel electrophoresis, purity using Nanodrop^TM^ 2000 (Thermo Fisher Scientific), and concentration using Qubit^®^ 2.0 (Thermo Fisher Scientific). To prepare the Illumina sequencing libraries, we used the NEBNext^®^ Ultra^TM^ FS II Library Prep kit (New England Biolabs) with default parameters. Approximately 500 ng of gDNA were used for each sample as input. Libraries of around 350 bp were generated after enzyme digestion of samples with high DNA quality. Herbarium samples were not digested. Quality control was evaluated using BioAnalyzer 2100 (Agilent), and concentration using Qubit^®^ 2.0. The enrichment step used baits specifically designed for Annonaceae by Daicel Arbor Biosciences merging the original Annonaceae kit (Couvreur et al., 2019), with genes selected from the Angiosperms353 panel (Johnson et al., 2019). With this strategy, we targeted exons from 799 nuclear genes. Samples were enriched in sets of 8–10 samples, combined based on the type of material (silica dried or herbarium samples), and concentration of the sample. The enrichment step lasted for 24 hours at 60° C, below the 65° C suggested by Daicel Arbor Biosciences (https://arborbiosci.com) to assure the baits would capture samples from *Asimina*. Initial trials using 62° C for 16 hours and 20 hours recovered extremely low concentrations of DNA and the parameters were updated. The final enriched sets of libraries were sequenced by Macrogen (https://macrogen-europe.com/finalized) using an Illumina NovaSeq 6000 platform. This platform generated around 2 Gb of data for each sample and paired-end reads of 150 bp.

### Sequence analyses

Illumina reads were demultiplexed at the sequencing facility and low-quality reads were trimmed using fastp 0.23.4 (Chen, 2023) with a quality phread score threshold of 30 (-q 30). Quality control was assessed using FastQC 0.12.1 (Andrews, 2010), and MultiQC (Ewels et al., 2016). HybPiper 2.1.5 (Johnson et al., 2016), as available in the Vlaams Supercomputer Centrum (VSC-HPC; https://www.vscentrum.be/), was used to assemble the genes. As input for the pipeline, we used the 799 genes selected as references (-t_dna), paired-end reads (-r), unpaired reads (--unpaired), the BWA 07.17 (Li and Durbin, 2009) aligner (--bwa), and ran the intronerate script (--run_intronerate) for up to 1200 seconds for each gene (--timeout_exonerate_contigs). The remaining parameters of HybPiper were left as default. We retrieved FASTA files of Exon-Only and Supercontig (exons + introns/intergenic-regions) sequences for all samples. The following HypPiper commands were run to obtain summary statistics; all assembled genes (retrieve_sequences); and paralog statistics (paralog_retriever). Two genes completely failed, and two were excluded due to low representation (i.e., they appeared in less than 80% of the samples). To evaluate the recovery of plastome genes, we used the pipeline HybPiper 2.1.5 (Johnson et al., 2016) with the same parameters as described before, and a reference file with sequences of 78 protein-coding genes from the plastome of *Annona muricata* L. (Niu et al., 2020; GenBank: MT742546.1). Attempts to assemble complete or nearly complete plastomes using the pipeline GetOrganelle 1.7.4.1 (Jin et al., 2020) resulted in fragmented and incomplete results, so here we will focus on the assemblies obtained using protein coding genes as references.

### Sequence alignments and statistics

Each step detailed here was repeated for the Exon-Only, Supercontig, and Plastome datasets. Gene sequences were aligned using MAFFT 7.450 (Katoh and Standley, 2013) with an automatic selection of alignment mode and a maximum of 1,000 interactions. Poorly aligned regions and bases with more than 50% of the samples as missing data were removed using ClipKIT (Steenwyk et al., 2020). Alignments were concatenated using AMAS (Borowiec, 2016) to generate a supermatrix with all molecular data combined. Statistical properties of each alignment, including the combined alignments, were obtained using the R (R Core Team, 2023) package *ape* (Paradis and Schliep, 2019). Parsimony informative sites (pis) were identified using the R package *ips* (Heibl, unpublished) for complete and *Asimina*-only alignments. The genetic distance between samples of *A. triloba* was also evaluated using the package *ape*, function “dist.dna,” and the K80 evolutionary model.

### Comparison between Custom and Original Angiosperms353 sets

To further evaluate the efficiency of the Custom Angiosperms353 bait kit, we compared the results obtained with data generated for all eleven species of *Asimina* using the Original Angiosperms353 kit during the enrichment step. Of the 11 specimens sampled in this new enrichment, three were shared between experiments (i.e., *A. rugelii*, *A. tetramera*, and *A. triloba*), and the remaining species were from the same or nearby populations (Table S2). The laboratory procedures and sequencing followed Couvreur et al. (2019). In short, DNA was extracted from silica-dried samples using the MATAB and chloroform separation methods (Mariac et al., 2014). Sequencing libraries were constructed following a modified protocol of Rohland and Reich (2012) using 6-bp barcodes and Illumina indexes to allow the multiplexing. After cleaning and quantification, libraries were combined in sets of eight. Targeted regions were enriched using the designed baits for Annonaceae, and amplified using 14–16 cycles. Libraries were sequenced on an Illumina HiSeq v3 platform using paired-end reads of 150 bp at CIRAD facilities (Montpellier, France). Gene assembly steps were the same as previously described.

## RESULTS

### Target sequences

We recovered an average of 17,368,883 paired-end reads after cleaning with fastp. The maximum number of reads was 27,276,263 (*A. reticulata*), and the minimum number of reads was 4,754,971 (*G. laoticus*). We obtained 49.5% (31.5–70.1) of on-target reads on average (Table 1). On average 798.4 genes were obtained per sample (99.9%), ranging from a minimum of 783 genes (*G. laoticus*) to a maximum of 802 genes (*A. manasota26*, *A. parviflora55*, and *A. reticulata39*). We assembled genes with at least 50% of size for 776.9 genes on average, ranging from a minimum of 748 genes (*G. laoticus*), and a maximum of 788 genes (*A. triloba49*) (Figure 1; Table 1). Four genes, two from the Angiosperms353 set and two from the Annonaceae set, completely failed or were present in less than half of the samples and were excluded (Figure 1). Ten genes recovered assemblies shorter than 200 bp and were excluded as well. Paralogs were flagged for 90 genes, 56 genes with a single specimen flagged, and 70 genes with less than five samples flagged. Of the 90 genes flagged, 16 were from the Angiosperms353 and 74 from the Annonaceae set. After removing putative paralogs, the final number of genes available to be used for phylogenetic studies of Annonaceae was 701.

**FIGURE 1.**
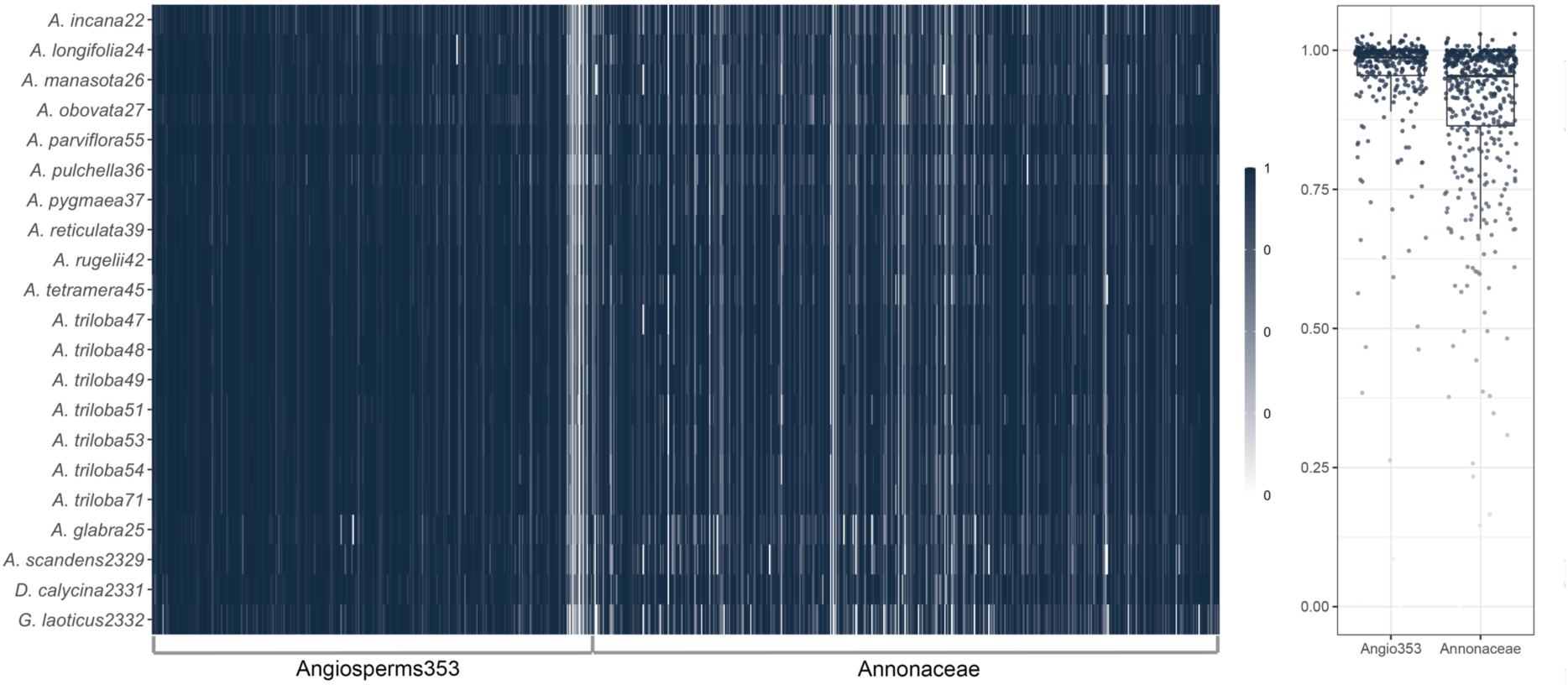
Proportion of gene recovery. A. Heatmap for all the 799 genes targeted. Each row corresponds to a different specimen, and each column corresponds to a different gene. Different shades of blue correspond to different proportions of gene assembly considering the reference. B. Boxplot with values with proportional values of gene assembly for genes from the Angiosperms353 panel, and genes from the Annonaceae panel.

**TABLE 1.**
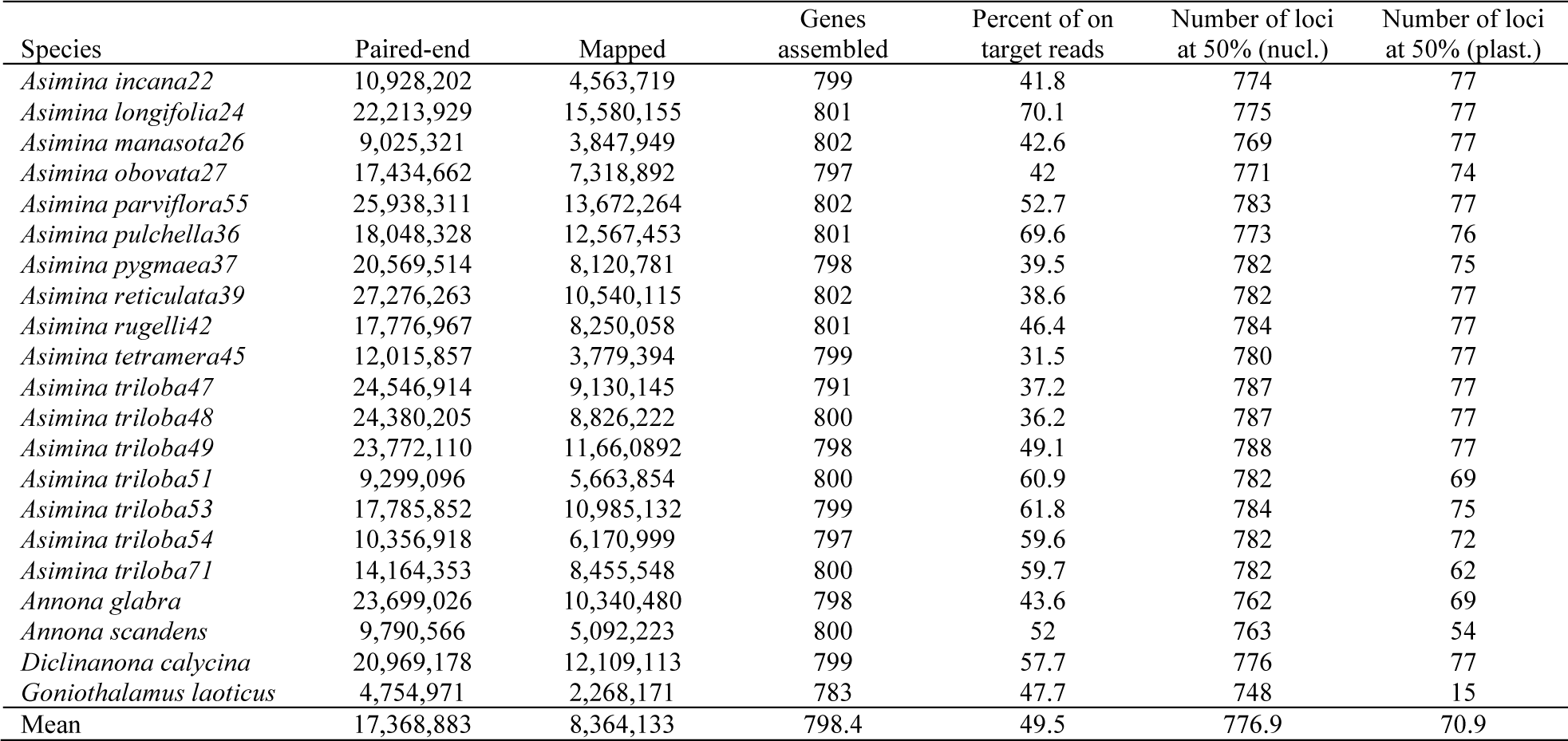
Summary statistics of sequencing success including the number of raw paired-end reads obtained, percentage of on-target reads, number of loci obtained, percentage of gene recovery, and number of loci retained after paralogs removed.

The length of the per-gene alignments ranged from 201 bp to 5,652 bp, with a mean value of 862.9 bp. Alignments from the Angiosperms353 set ranged from 240 bp to 4,653 bp, with a mean size of 984.5 bp. The extreme values were obtained from the Annonaceae set, with a mean value of 763.2 bp. The number of pis for “complete” and “*Asimina*-only” alignments ranged from zero/zero to 360/122, with a mean value of 49.1/13.5 sites, respectively (Figure 2). The number of pis from the Angiosperms353 set ranged from zero/zero to 232/122, with a mean value of 56/15.8 sites, respectively. The number of pis from the Annonaceae set ranged from one/zero to 360/96, with a mean value of 43.5/11.5 sites (Figure 2). The total length of all per-gene alignments combined (Exon-Only) was 604,924 bp, of which 42,924 were pis when outgroups were considered, and 14,037 were pis when only species of *Asimina* were considered (Table 2). The Angiosperms353 alignment was 311,095 bp long, of which 18,426 were pis when outgroups were considered, and 5,735 were pis when only species of *Asimina* were considered. The Annonaceae alignment was 293,829 bp, of which 24,498 were pis when outgroups were considered, and 8,302 were pis for *Asimina* only (Table 2). The pairwise genetic distance between samples of *A. triloba* ranged from 0.13% to 0.326% (Table 3).

**FIGURE 2.**
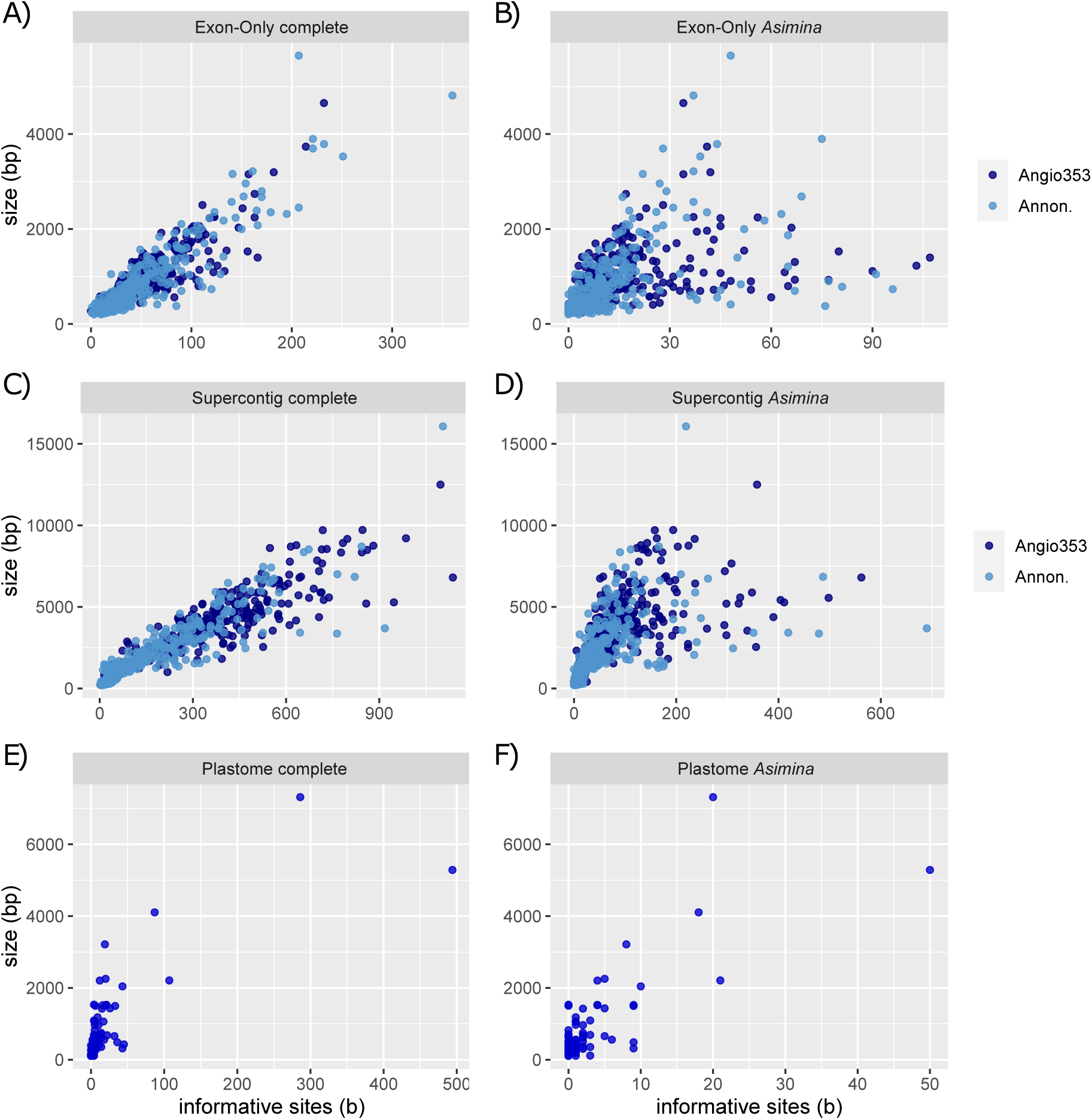
Size and number of parsimony informative sites for each of the 701 genes compared. In dark blue the genes from the Angiosperms353 panel; in steel blue the genes from the plastome; and in light blue the genes from the Annonaceae panel. A–B. Genes from Exon-Only dataset, with (A) and without outgroups (B). C–D. Genes from Supercontig dataset, with (C) and without outgroups (D). Genes from Plastome dataset, with (E) and without outgroups (F).

**TABLE 2.** Values of aligned sequences and parsimony informative sites for each dataset.

|  | Sites | Informative Sites |  |
| --- | --- | --- | --- |
|  |  | Complete | *Asimina* only |
| Angiosperms353 | 311,095 | 18,426 | 5,735 |
| Annonaceae | 293,829 | 24,498 | 8,302 |
| Exon-Only | 604,924 | 42,924 | 14,037 |
| Angiosperms353 | 1,198,434 | 348,929 | 261,570 |
| Annonaceae | 740,649 | 175,598 | 128,239 |
| Supercontig | 1,939,083 | 524,527 | 389,809 |
| Plastome | 66,237 | 8,092 | 1250 |

**TABLE 3.** Comparison of within species molecular variation for eight samples of *Asimina triloba* using the K80 model. Values above the diagonal are pairwise comparisons for the Exon-Only dataset, and values below the diagonal are pairwise comparisons for the Supercontig dataset.

|  | *A. tri47* | *A. tri48* | *A. tri49* | *A. tri51* | *A. tri53* | *A. tri54* | *A. tri71* |
| --- | --- | --- | --- | --- | --- | --- | --- |
| *A. triloba*47 | 0 | 0.002349 | 0.002201 | 0.002770 | 0.003157 | 0.003240 | 0.003101 |
| *A. triloba*48 | 0.003098 | 0 | 0.001966 | 0.002430 | 0.003084 | 0.003257 | 0.003014 |
| *A. triloba*49 | 0.003176 | 0.003402 | 0 | 0.002507 | 0.003084 | 0.002186 | 0.002719 |
| *A. triloba*51 | 0.004655 | 0.004379 | 0.004868 | 0 | 0.003055 | 0.003265 | 0.002970 |
| *A. triloba*53 | 0.003815 | 0.004009 | 0.002729 | 0.004796 | 0 | 0.001301 | 0.002181 |
| *A. triloba*54 | 0.004147 | 0.004227 | 0.002922 | 0.005269 | 0.002607 | 0 | 0.002213 |
| *A. triloba*71 | 0.004048 | 0.003792 | 0.003799 | 0.004580 | 0.003552 | 0.004067 | 0 |

### Nuclear non-targeted sequences

Off-target regions from introns and intergenic regions collectively known as “splash-zone” in target-enrichment studies, were obtained for *Asimina* and outgroups. In total, off-target regions were obtained for 499 genes, 277 genes from the Angiosperms353 set and 222 from the Annonaceae set. The length of the alignments ranged from 656 bp to 16,069 bp, with a mean value of 3,677 bp. Alignments from the Angiosperms353 set ranged from 824 bp to 12,498 bp, with a mean size of 4,222 bp. The extreme values were again obtained from the Annonaceae set, with a mean value of 2,997.1 bp. The number of pis for “complete” and “*Asimina*-only” alignments ranged from 10/three to 1,137/690, with a mean value of 331.4/120 sites, respectively (Figure 2). The number of pis from the Angiosperms353 set ranged from 14/5 to 1,137/562, with a mean value of 392.1/142.4 sites, respectively. The number of pis from the Annonaceae set ranged from 10/3 to 1,105/690, with a mean value of 255.5/92.1 sites (Figure 2). When all the alignments were combined (Supercontig), the length was 1,939,083 bp when including the 701 genes. When outgroups were considered, the data set contained 524,527 pis, and 389,809 pis in the“*Asimina*-only” alignments (Table 2). When the different datasets are evaluated, the Angiosperms353 alignment was 1,198,434 bp long, of which 348,929 were pis when outgroups were considered, and 261,570 were pis when only species of *Asimina* were considered; while the Annonaceae alignment was 740,649 bp long, of which 175,598 were pis when outgroups were considered, and 128,239 were pis for *Asimina* only (Table 2). The pairwise genetic distance between samples of *A. triloba* ranged from 0.26% to 0.52% (Table 3).

### Plastome non-targeted sequences

Off-target regions from the plastome were obtained for *Asimina* and outgroup. In total, off-target sequences were obtained for all 78 genes used as references (Figure 3), of which *pet*L, *pet*N, and *rpo*A were excluded because they had sequences for less than half of the individuals or had sequences shorter than 100 bp. The retained sequence alignments ranged from 105 bp to 7,314 bp, with a mean value of 883.2 bp (Figure 1; Table 1). The number of pis for “complete” and “*Asimina*-only” alignments ranged from zero/zero to 494/50, with a mean value of 21.9/3.2 sites, respectively (Figure 2). The length of all the alignments combined (Plastome) was 66,237 bp, of which 8,092 bp were pis when outgroups were considered, and 1,250 bp were pis when only species of *Asimina* were considered (Table 2).

**FIGURE 3.**
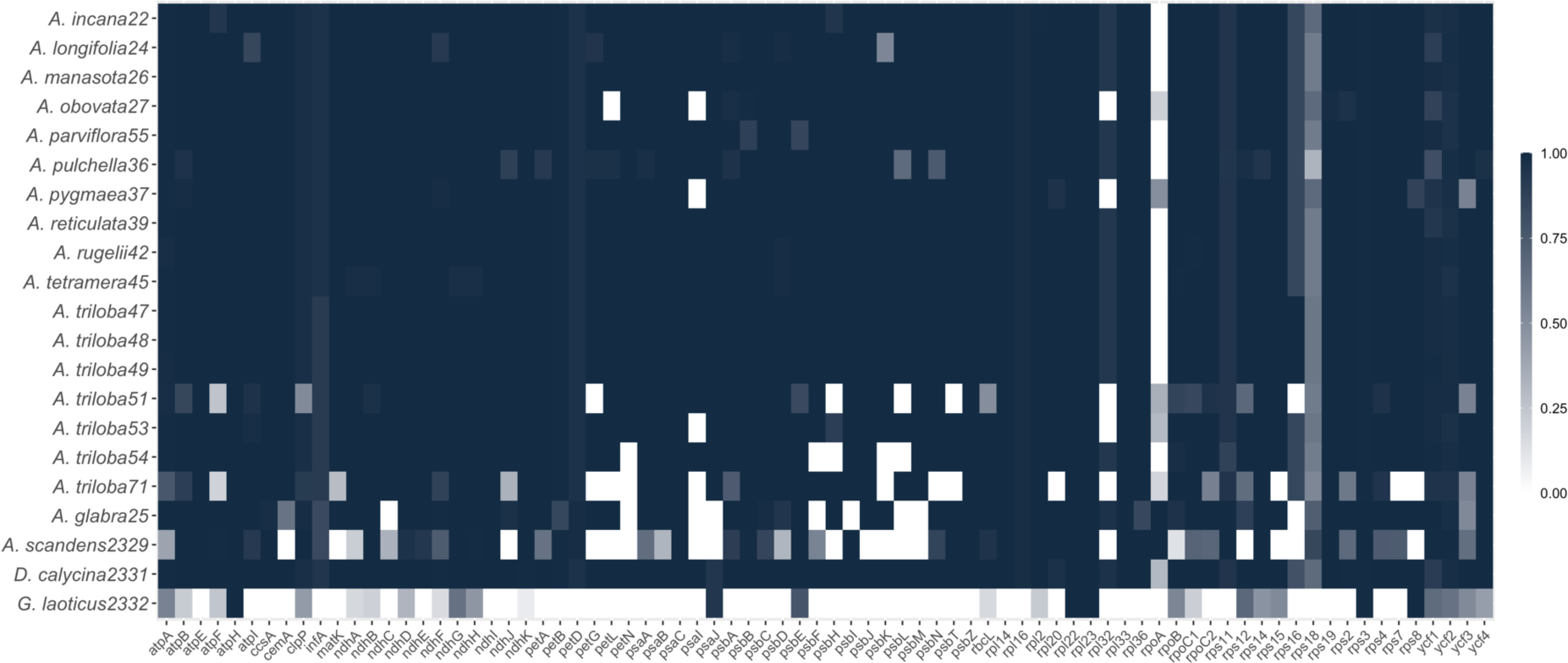
Heatmap for all the 78 genes from the plastome. Each row corresponds to a different specimen, and each column corresponds to a different gene. Different shades of blue correspond to different proportions of gene assembly considering the reference.

### Comparison between Angiosperms353 baiting kits

The Custom Angiosperms353 bait kit tackled 334 genes, 332 of which could be assembled, while the Original kit tackled all the 353 genes, 350 of which could be assembled for *Asimina* species (Table 4). The mean value of gene recovery for the Custom kit was 0.94, while the value for the Original bait kit was 0.79 (Figure 4; Table 4). The Custom genes assembled had a longer combined alignment of 328,598 bp, compared to the Original genes with a combined alignment of 265,911 bp. When the number of pis are compared, the pattern is reversed, and more variable sites are observed in the Original bait kit (5,145 vs. 3,878 bp). For Supercontig regions, the Custom kit recovered 285 and the Original 216 genes. The assembled Custom genes had a longer combined alignment of 1,693,166 bp, compared to the Original genes with a combined alignment of 1,133, 674 bp. The Custom set recovered 231,718 bp of pis and the Original kit 126,163 bp of pis (Table 4).

**FIGURE 4.**
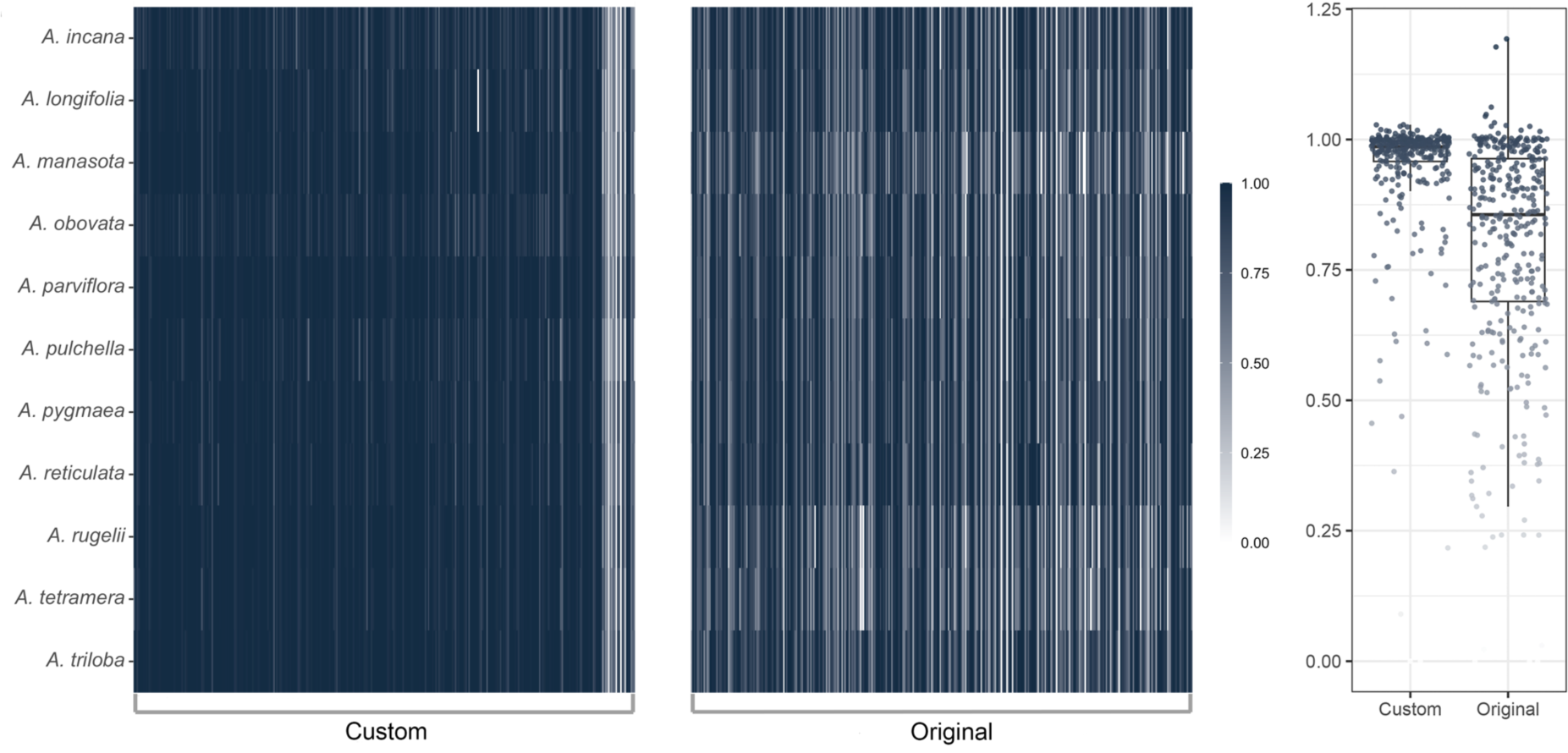
Proportion of gene recovery. A. Heatmap comparing the results of Custom and Original Angiosperms353 gene set. Each row corresponds to a different specimen, and each column corresponds to a different gene. Different shades of blue correspond to different proportions of gene assembly considering the reference. B. Boxplot with values with proportional values of gene assembly for genes of the Custom and Original Angiosperms353 sets.

**TABLE 4.**
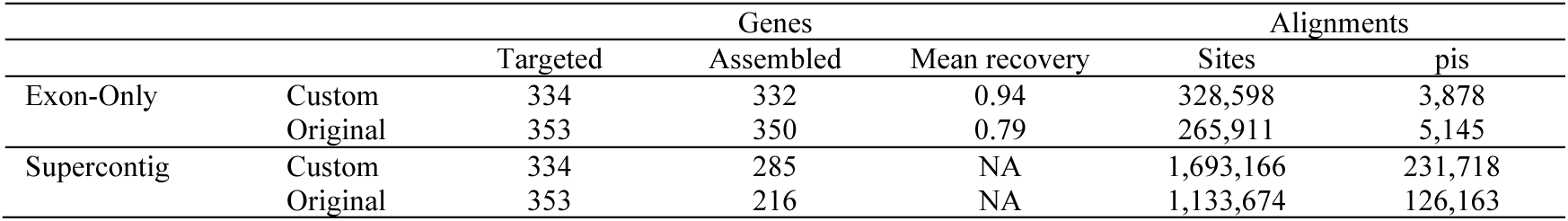
Comparison between the Custom and Original Angiosperms353 bait kits using one individual of each species of *Asimina*.

|  |  | Genes |  |  | Alignments |  |
| --- | --- | --- | --- | --- | --- | --- |
|  |  | Targeted | Assembled | Mean recovery | Sites | pis |
| Exon-Only | Custom | 334 | 332 | 0.94 | 328,598 | 3,878 |
|  | Original | 353 | 350 | 0.79 | 265,911 | 5,145 |
| Supercontig | Custom | 334 | 285 | NA | 1,693,166 | 231,718 |
|  | Original | 353 | 216 | NA | 1,133,674 | 126,163 |

## DISCUSSION

Hybridization capture-based technologies enable the retrieval of hundreds of nuclear loci from diverse plant lineages (e.g., McKain et al., 2018; Johnson et al., 2019). The specificity of custom bait kits, and the various parameters used during the enrichment step can assure high levels of recovery for the targeted regions (McKain et al., 2018). Here, we provide an updated version of the Annonaceae bait kit (from now on Annonaceae799), combining the Angiosperms353 kit, with the original Annonaceae kit and targeting 799 genes. The efficiency of this new kit is extremely high with only 2 genes completely failing, a mean value of 776.9 genes assembled at 50% per species. The assembly of off-target regions was also high, with 499 genes obtaining sequences from the “splash-zone” (i.e., introns, and intergenic regions). Of these, 277 genes were targeted by the Angiosperms353 panel. The enrichment step implemented also allowed the assembly of non-nuclear off-target regions, such as 78 protein coding genes from the plastome, allowing the use of different genomic compartments for comparative and evolutionary studies (Weitemier et al., 2014).

### Target enrichment in plant studies

In order to make the original Annonaceae baiting kit more universal, especially with the prospect of the Plant and Fungus Tree of Life project, we generated an augmented version of the Annonaceae baiting kit (Couvreur et al., 2019) by including the Angiosperms353 exons. This new kit allows the sequencing of 799 single to low-copy nuclear genes, instead of the original 469 exons from the original kit. The new version of the kit does not imply an increase in costs, since it uses the maximum number of baits available by Daicel Arbor Bioscience entry-level kits (20K). The advent of hybrid kits, including baits designed using genes selected from universal kits (i.e., Angiosperms353), and family specific kits, allows the interconnection of data generated at broad taxonomic scales with projects focusing on lower taxonomic scales (Baker et al., 2021). The number of genes assembled by using universal kits is one of its limitations, since the number of baits to be used is a bottleneck in any kit and it is challenging to embrace all the molecular variation present within the flowering plants. The solution to maximize molecular variation while keeping the number of different baits to a minimum reached its limitation for some species and clades, as was clear in the original publication (Johnson et al., 2019). The maintenance of the Angiosperms353 gene set coupled with clade specific baits is a step forward assuring high recovery values for all the genes in most cases (e.g., Ogutcen et al., 2021; Fonseca et al., 2023). The recovery rate of the new version of the Annonaceae bait kit is extremely high (Figure 1), with a mean value of 798.4 genes or 99.9%, and 776.9 genes assembled at up to 50% per species (Table 1), echoing previous results obtained for Bignoniaceae of 98% (Fonseca et al., 2023).

The merger of gene sets provides the best of both worlds: the connection between projects with different taxonomic scopes, and the inclusion of high variable genes to resolve shallow relationships in the tree of life (Hendricks et al., 2021). Positive results were recently obtained for diverse plant clades that implemented this approach (Ogutcen et al., 2021; Fonseca et al., 2023), but the second assertion was not always met. Some genes from the Angiosperms353 were as good as genes from lineage-specific bait kits (e.g., Larridon et al., 2019), in some cases even outperforming the latter in terms of variation and number of pis (Fonseca et al., 2023). This is also true for the Annonaceae799 kit, with values of variation and pis for the genes from the Angiosperms353 panel outperforming the genes from original Annonaceae kit (Figures 2; Table 2). The genes from the Angiosperms353 were also longer and recovered more regions from the “splash-zone,” when compared to the original Annonaceae bait kit. These findings are the result of the gene selection process (Johnson et al., 209; Leebens-Mack et al., 2019), favoring longer genes with more introns compared to the Annonaceae panel (Couvreur et al., 2019).

We obtained 49.5% of on-target reads (Table 1), a value comparable to other flowering plant clades (e.g., 31.6% in *Dioscorea*, Soto Gomez et al., 2019; 48.6% in *Euphorbia*, Villaverde et al., 2018). This relatively high number of off-target reads allows the assembly of plastome genes through the HybPiper (Johnson et al., 2016). The recovery of off-target regions after target-enrichment reactions has been long suggested, and even the name Hyb-Seq was coined for this protocol (Weitemier et al., 2014). It is not clear how to implement the necessary parameters for the enrichment step necessary to assure high recovery rates for on and off-target regions (Weitemier et al., 2014). In fact, some studies failed to assemble complete/nearly-complete plastomes or plastome genes for most of the samples (e.g., Villaverde et al., 2019). Here we followed the protocol provided by Daicel Arbor Biosciences 5.02 (https://arborbiosci.com), with an enrichment step lasting for 24 hours at 60° C. The temperature is notably lower than 65° C suggested by Daicel Arbor Biosciences. However, for the new version of the Annonaceae bait kit it guarantees extremely high rates of recovery for on-target, and off-target regions combined (Figures 1, 4).

### Annonaceae799 baiting kit

The new version of the Annonaceae bait kit will allow users to infer phylogenies, study species complexes, and infer population histories using exons from up to 799 genes. Considering the *Asimina* framework with four outgroups (viz., *Annona glabra*, *Annona scandens*, *Diclinanona calycina*, and *Goniothalamus laoticus*), 90 genes were flagged as containing paralogs, leading to 709 single copy genes. Of the 90 genes flagged, 16 are from the Angiosperms353 and 74 from the Annonaceae set. Considering previous studies (Bignoniaceae, with 202 out of 711 genes flagged, Fonseca et al., 2023; Gesneriaceae, with 219 out of 830, Ogutcen et al., 2021), the number of genes flagged here is relatively low. It is clear that the paralogs are minor problem if we consider that 56 genes had a single specimen flagged, and 70 genes had less than five samples flagged. Further analyses using tree methods (Yang and Smith, 2014; Morales-Briones et al., 2021) can also increase the number of useful genes, and even provide new and hidden orthologous sets among the current reference genes (Yang and Smith, 2014). However, it is relevant to recognize that the number of paralogs is always dependent of the phylogenetic context as well as the sequencing depth (Johnson et al., 2016), so the values observed here should be considered with caution and more comprehensive studies on Annonaceae will probably find more paralogs.

The original Annonaceae baiting kit was designed to be used from the large scale family level all the way down to within-population phylogeography, and has been very successful (Lopes et al., in press; Helmstetter et al., 2020a; Helmstetter et al., 2020b; Dagallier et al. 2023; Martínez-Velarde et al. 2023). The new set designed here, was first intended to tackle taxonomically complex taxa, as well as population studies using the maximum amount of molecular data possible through target-enrichment (up to 20K baits). This is the case for the genus *Asimina*, composed of putative ten species and many accounts of possible gene flow (and introgression) between species and the formation of hybrids (Kral, 1960). The first application of the Annonaceae799 baiting kit will be in a phylogenomic and population study of the genus *Asimina* (Fonseca et al., in prep.). The molecular variation within the genus is low compared to the results including outgroups (Figure 2; Table 2). However, thousands of bases are available for species-level phylogenetic relationships of *Asimina* (Figure 2; Table 2), and hundreds of bases when different individuals of *A. triloba* are compared (Table 3), suggesting that enough within-species molecular variants for population studies are available (Casillas and Barbadilla, 2017). A positive surprise is the performance of the genes from the Angiosperms353 panel, compared to the original Annonaceae bait kit in terms of molecular variation (Figure 2; Table 2). The new genes certainly improved the original bait kit beyond the increase in the number of bases and genes, providing more variable bases and pis for phylogenetic and populational studies. The new version of the Annonaceae bait kit will also allow for data reuse and will contribute to ongoing efforts to reconstruct the plant tree of life using the Angiosperms353 kit (Baker et al., 2021) as a source of molecular data.

### Angiosperms353 gene sets

Here we provided a comparison between two different bait kits targeting the same set of genes. The Original Angiosperms353 bait kit was designed to target the same genes for all flowering plants and included a comprehensive set of more than 75,000 different baits for this objective (Johnson et al., 2019). To reduce the amount of reads necessary to achieve the same objective in a smaller clade of the flowering plants, we used the 410 alignments from Leebens-Mack et al. (2019) and kept only the genes from Angiosperms353 set (Johnson et al., 2019), and the species of Annonaceae (i.e., *Annona muricata* and *Uvaria macrocarpa*). With this strategy we reduced the number of necessary baits to target a subset of the genes (334 out of the 353) and use a fraction of the baits (just 9.3%). Despite targeting fewer genes, the Custom bait kit recovered longer combined alignments for both Exon-Only and Supercontig. The number of pis varied and were higher for the Original kit when Exon-Only genes were compared, and lower for the same kit when Supercontig genes were compared (Table 4). These results reveal the editions made by Johnson et al. (2019) in the original 410 alignments from the Leebens-Mack et al. (2019). Low variable exons were removed, while more variable exons were probably added, leading to the discrepancies observed when Exon-Only results are compared. The maintenance of low variable exons shows its importance when Supercontig results are compared (Table 4). The size of the combined alignment of Supercontig is almost 50% larger and includes 84% more pis, providing more informative positions. Highlights of the comparison between strategies to obtain sequences for the same genes are: (1) the higher efficiency of the Custom kit during gene assembly compared to the Original Angiosperms353, as already suggested by Baker et al. (2021); (2) The higher number of pis for the Custom kit, allowing more positions to be used during tree search at shallow levels of the phylogeny. The latter observation is extremely important for individuals and research groups designing custom bait kits for shallow levels of the angiosperms tree of life, or in intra-specific studies and planning to use the Angiosperms353 gene set.

Here we showed that the inclusion of the Angiosperms353 gene set in a custom baiting kit can save thousands of baits, while recovering better results compared to the original kit. Depending on the references used for the Angiosperms353 gene set (i.e., the original alignments from Leebens-Mack et al., 2019), it is also possible to recover larger assemblies from the same genes as shown here. As was clear, combining kits is possible and could be technically superior. The final choice on including the Angiosperms353 gene set in the researcher’s custom kit or use different kits separated, however, also depends on other factors such as technical or financial limitations of sequencing many hundreds of genes or the pricing policies of the companies responsible for synthesizing the baits and making available the enrichment kits.

## AUTHOR CONTRIBUTIONS

L.H.M.F, T.L.P.C., and L.W.C. conceived the study; L.H.M.F and L.W.C wrote the manuscript; L.H.M.F., V.S., and F.J.N. obtained and analyzed the data; P.A. and V.S. assisted with laboratory work including DNA extractions, library preparations, and the enrichment step, and provided feedback on the manuscript; K.R.G., K.M., and F.A.M. provided samples and feedback on the manuscript. All authors approved the final version of the manuscript.

## ACKNOWLEDGEMENTS

L.H.M.F and L.W.C would like to thank UGent and Foundation Arboretum Wespelaar for funding the project. This project also received funding from the European Research Council (ERC) under the European Union’s Horizon 2020 research and innovation program (grant agreement No. 865787). K.M. would like to thank Maxwell-Hanrahan Foundation for supporting field work. The authors are thankful to Dr. Anne Cox, and Dr. Élaine Norman for help during fieldwork and extremely valuable information about species locality; Dr. Robert Raguso and Dr. Deise Gonçalves for sharing silica-dried samples; and Dr. Lucas Majure, and Kent Perkins for sharing silica dried and leaf fragments from specimens housed in FLAS herbarium. We would also like to thank the Vlaams Supercomputer Center (VSC) at Ghent University for the computational infrastructure made available and all the guidance and support.

## DATA AVAILABILITY STATEMENT

Raw sequencing reads are available via the National Center for Biotechnology Information (BioProject: XXX); custom scripts are available at GitHub (https://github.com/luizhhziul/Asimina).

